# RNA trans-splicing treatment for Duchenne muscular dystrophy

**DOI:** 10.64898/2026.09.13.751281

**Authors:** Ryan H. Hsu, Claire E. Williams, Geraldine Maier, Miriam Gullo, Karen Lettieri, Cindy J. Alvarez, Kip J. Hermann, Stella Kramer, Satchidananda Panda, Lukas C. Bachmann, Samuel L. Pfaff

## Abstract

Duchenne muscular dystrophy (DMD) is a fatal disorder caused by *dystrophin* mutations, leading to progressive muscle degeneration and cardiac failure. Although AAV-based DMD therapies with innovative designs to overcome the cargo limits of the virus are effective in animal models, these current systems may face clinical translation challenges related to efficiency, off-target effects, and immunogenicity. We developed a multi-vector RNA End Joining (REJ) system to split large genes into multiple co-delivered AAVs that engage the cell’s intrinsic spliceosome to precisely reassemble RNA segments that encode very large scar-free proteins. We show the system is efficient and has negligible off-target interactions. *In vivo*, REJ vectors expressing the adenine base editor Abe8e, mini-dystrophin Dp253, or native full-length dystrophin Dp427 each prevented muscle degeneration. Machine-learning histopathology of more than 120,000 myofibers revealed robust reductions in nuclear infiltration, improved centronucleation, and normalized hypertrophy, supported by functional, transcriptomic, and behavioral analyses. These findings establish REJ as a clinically viable AAV-based strategy for DMD that enables expression of large therapeutic proteins while avoiding potentially antigenic bacterial protein- or DNA-recombinases.

## Main

Duchenne muscular dystrophy (DMD) is a fatal X-linked disorder caused by mutations in the *dmd* gene and affects approximately 1 in 5,000 male births^1–3^. These mutations eliminate functional dystrophin, a 427 kDa protein encoded by an 11.1 kb transcript, disrupting the mechanical link between the actin cytoskeleton and membrane-associated glycoproteins in skeletal and cardiac muscle. Without this structural support, muscle fibers undergo repeated injury and progressive degeneration. Symptoms usually emerge at approximately three years of age, followed by loss of ambulation and ultimately fatal respiratory or cardiac failure by about 30 years of age^1,2^. Preclinical studies have shown that restoring dystrophin can improve disease outcomes^4–13^, making DMD a strong candidate for gene therapy^14^. However, the limited 4.7 kb cargo capacity of adeno-associated virus (AAV) vectors remains a major barrier (Figure 1A)^4,15–17^. Although full-length dystrophin (Dp427) may offer the greatest therapeutic benefit, its coding sequence is far too large for a single AAV. We therefore sought to develop an RNA trans-splicing strategy capable of expressing large therapeutic proteins in vivo.

**Figure 1.**
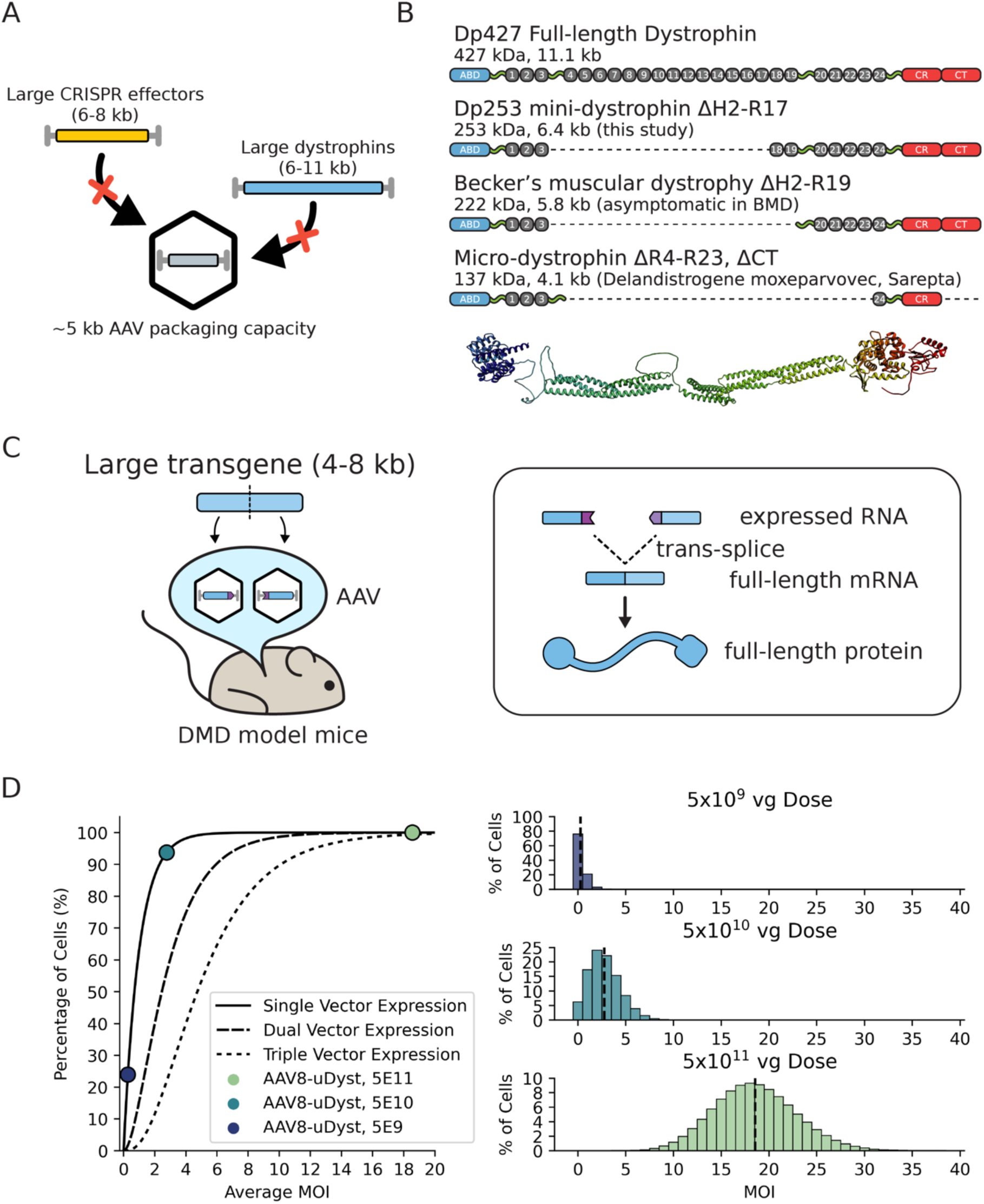
Multi-vector strategies for Duchenne muscular dystrophy. A) The 4.7 kb cargo capacity of AAV precludes the transfer of large therapeutic proteins such as some CRISPR effectors and the native 427 kDa dystrophin protein. B) Engineered variants of dystrophin contain varying domains of the native protein. The ΔH2-R17 mini-dystrophin Dp253 (this study) contains the full N- and C-terminal domains and the R18-R19 domains absent in some asymptomatic ΔH2-R19 BMD patients. Predicted protein structure of the ABD, hinges, rods, and CT domains of a representative micro-dystrophin shown (below). C) A large coding sequence (blue) is divided into two segments, each delivered by a separate AAV and appended with a short REJ module (purple) containing splice enhancing elements, splice donor/acceptor motifs, and RNA dimerization domains. RNA dimerization of the two transcribed RNAs into a pseudo-linear substrate positions a splice donor on the 5′ RNA and a splice acceptor on the 3′ RNA. RNA end-joining (REJ) occurs when the endogenous spliceosome recognizes this configuration as a conventional intron and excises it, joining the two segments into a single continuous mRNA that encodes full-length, scar-free protein. D) Stochastic modeling of multi-vector AAV transduction efficiency. A Poisson process model was used to calculate the effective multiplicity of infection (MOI) distribution across viral doses using a micro-dystrophin vector. The model indicates that high MOIs are achievable within clinical dosing ranges. For 90% cellular co-expression, a dual and triple vector system requires 2.6-fold and 4.4 fold increase in viral dose, respectively.

Current AAV therapies largely rely on micro-dystrophins, such as Dp137, that are sufficiently small for delivery in one vector^5–7,18,19^. However, these shortened proteins lack domains required for optimal muscle stability and have demonstrated limited efficacy in clinical trials (Figure 1B)^20,21^. Studies of Becker muscular dystrophy suggest that larger, intermediate dystrophins, including Dp253, retain substantially greater functional activity^5,22,23^, but these constructs also exceed the capacity of a single AAV. Multi-vector strategies can divide large genes among separate vectors and reassemble them through DNA-, RNA-, or protein-level mechanisms^8–10,15,16,24–27^. Nevertheless, DNA recombination is often inefficient and imprecise, requiring high viral doses^9,28^, whereas bacterially derived inteins may provoke immune responses, leave residual peptide scars, and produce potentially disruptive protein fragments^29^. Newer systems, including StitchR and AAVLINK, allow improved large-gene delivery^13,30^, but limitations in efficiency and antigenicity remain. Similar packaging constraints affect CRISPR-based therapies because adenine base editors such as Abe8e, as well as many advanced editors with improved specificity, are also too large for a single AAV^11,31–33^.

To overcome these limitations, we developed RNA End-Joining (REJ), an RNA trans-splicing platform that divides long coding sequences into AAV-compatible segments and appends short REJ modules to direct their precise assembly by the endogenous spliceosome (Figure 1C; Bachmann et al., companion manuscript, bioRxiv 2026). REJ generates full-length, scar-free mRNAs without foreign recombinases or protein-fusion elements and exhibits minimal off-target interactions. We used REJ vectors in DMD mouse models to deliver Abe8e for *dmd* gene repair or to express either Dp253 or native full-length Dp427 dystrophin (Figure 2A, E). Treatment was evaluated using molecular, transcriptomic, behavioral, physiological, and functional assays, together with machine-learning histopathology that quantified individual myofibers across whole-muscle sections. All three therapeutic approaches prevented major dystrophic features and preserved muscle function thereby establishing REJ as a potential clinically applicable platform for treating DMD. Collectively our observations treating DMD with multiple types of REJ-vectors support the possibility that this system may serve as a reliable platform for delivering other large therapeutic proteins with AAV while avoiding the inefficiency, protein scarring, and antigenicity associated with existing multi-vector systems.

**Figure 2.**
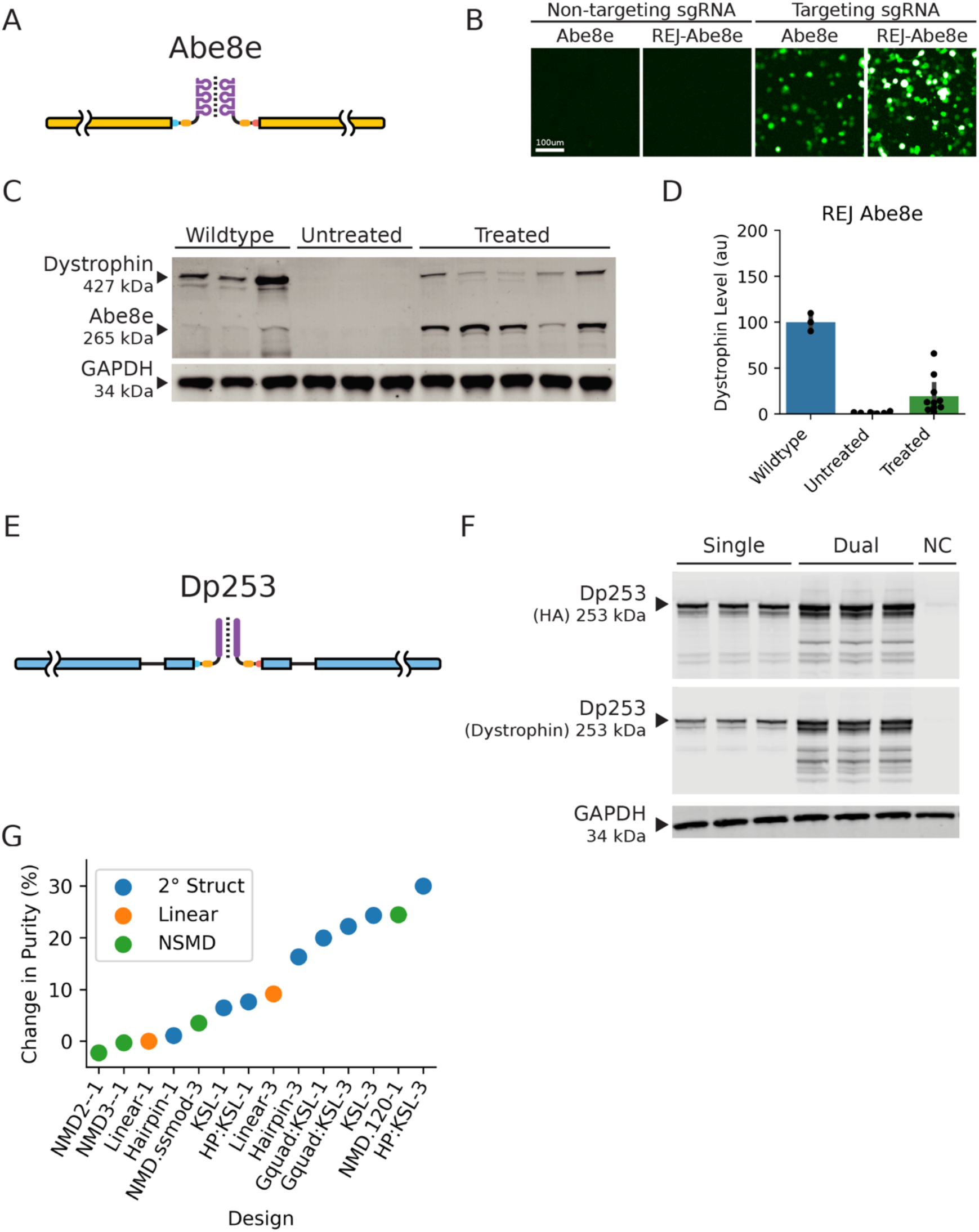
REJ vectors efficiently express Abe8e and Dp253. A) Adenine base editor Abe8e was split into two segments with appended REJ modules and co-delivered into cells using dual AAVs. The Abe8e coding sequence (yellow) was split into two trans-splicing exons with kissing stem loop RNA dimerization domains (purple) and the REJ intron modules, containing splice acceptor and donors (cyan, pink) and splice enhancers (yellow). Not shown, the AAVs additionally carry three loci for expression of sgRNA. B) HEK293T cells were transfected with a YFP reporter containing a region harboring the mdx.4cv nonsense mutation. Cells were additionally transfected with either a plasmid encoding full-length Abe8e or dual-plasmids encoding the split Abe8e, and either a targeting or non-targeting sgRNA. Cells transfected with dual Abe8e exhibited similar induction of YFP expression to those with the full-length control. C) Dual AAVs encoding split Abe8e were transduced into DMD mouse TA muscles. Abe8e was robustly expressed, resulting in restoration of endogenous Dp427 dystrophin expression. D) Abe8e-treated mice express 19.3% of wildtype dystrophin levels compared to 0.46% in untreated mice (p=0.0160). E) Dp253 (blue) was split into two trans-splicing exons with appended REJ modules (see A). An additional short stimulatory intron for enhanced protein translation was added to the 5’ and 3’ Dp253 RNA segments. F) HEK293T cells transfected with dual plasmids encoding Dp253 expressed high protein levels. G) Optimization of protein product purity. To prevent the translation of truncated protein fragments from un-spliced 3’ RNA, we screened 14 translation inhibition designs. While mechanisms such as nonsense mediated decay and G-quadruplexes showed promise, the most potent increase in purity levels (30%) was achieved by engineering a leading hairpin and using 3x kissing stem loops as the RNA dimerization domain in the 5’ UTR, coupled with a 3 times stoichiometric excess of the 5’ RNA vector.

## Results

### Establishing a viral dosing framework for multi-part AAV systems

Multi-vector gene therapy requires co-infection of target cells with complementary AAV vectors. Because AAV does not prevent superinfection, multiple vectors can deliver separate cargo segments to the same cell; however, the relationship between dose and co-infection efficiency is not well defined. We developed a stochastic Poisson model to predict co-expression from discrete AAV infection events (Figure 1D). To establish baseline transduction, mouse tibialis anterior muscles were treated with a single AAV encoding Dp153 across a range of doses, and dystrophin-positive myofibers were quantified by immunohistochemistry^18^. Fitting the model to these data showed that 90% of myofibers were transduced at 4.18 × 10^10^ vector genomes per muscle. Using empirically derived parameters from this titration, we modeled the co-infection needed for efficient reconstitution of split genes. Achieving 90% expression required a 2.6-fold higher dose for dual-vector systems and a 4.4-fold higher dose for triple-vector systems than for single-vector delivery. Although multi-vector approaches require increased dosing, predicted levels remain within established clinical safety parameters, supporting their feasibility for therapeutic gene delivery.

### Expression of gene editor Abe8e and mini-dystrophin Dp253

Having established the theoretical framework for multi-AAV dosing, we next designed REJ-based dual-vector strategies that could restore dystrophin expression in *dmd* mutant mice. Recent developments of novel CRISPR-Cas9 genome editors have demonstrated enormous potential in treating genetic disorders, but many of these systems use oversized fusion proteins and remain difficult to deliver. We designed a REJ dual-vector platform to express Abe8e, a large CRISPR-Cas9 adenine base editor (Figure 2A). This editor was expressed as two separate RNAs with REJ modules to induce RNA trans-splicing (Bachmann et al., companion manuscript, bioRxiv 2026), accompanied by three copies of a U6 promoter-driven sgRNA that targeted the *dmd^mdx.4cv^* mutation to restore endogenous dystrophin expression^34^.

To validate functional activity, dual REJ-Abe8e vectors or a single full-length Abe8e control plasmid were transfected into HEK293T cells. Together with these Abe8e vectors, either targeting or non-targeting sgRNA were included with a YFP reporter interrupted by a sequence containing the mdx.4cv nonsense mutation. Both the full-length control Abe8e and REJ-Abe8e constructs induced YFP expression when delivered with a targeting guide RNA, demonstrating efficient REJ-mediated Abe8e expression and functional editing activity (Figure 2B).

In parallel, we designed a dual-vector system to express the 253 kDa ΔH2-R17 mini-dystrophin based on those found in asymptomatic BMD (Dp253, Figure 2E). This REJ-Dp253 vector importantly includes the R18-R21 region protective for dilated cardiomyopathy and stimulatory introns to increase protein expression (Figure 2E). Western blotting of transfected HEK293T cells confirmed REJ-Dp253 expression at 148% of the single-plasmid Dp253 control, possibly due to more efficient transfection of the smaller split plasmids (Figure 2F).

The 3’ REJ-Dp253 RNA which encodes a potentially toxic C-terminal fragment of dystrophin was designed with a decoy out-of-frame start codon to prevent expression of the truncated protein fragment from unspliced RNA. Despite this, we noticed low but detectable levels of the C-terminal truncated protein, motivating us to develop mechanisms to prevent fragment expression. We designed and screened 14 experimental variations of the 3’ REJ RNA which contain a variety of translation inhibition mechanisms. These designs exploit the conditional presence of the 5’ UTR on the 3’ RNA, which is intact if unspliced, but removed when trans-spliced. We employed mechanisms including decoy reading frames to induce nonsense mediated decay, secondary structures to inhibit ribosome assembly or scanning, each individually demonstrating ability to improve purity of the full-length Dp253 (Figure 2G). In addition, we varied the transfection ratios of the 5’ and 3’ REJ RNA. Our screen demonstrates that modular inclusion of a leading hairpin structure and 3x kissing stem loops, delivered at a 3:1 5’ to 3’ ratio potently reduces fragment levels by 30% (Figure 2G).

Taken together, these *in vitro* studies revealed robust Abe8e activity and Dp253 expression using RNA trans-splicing to express the split genes.

### Dystrophin restoration and transcriptomic profile improvement in dystrophic muscle

To determine if REJ-enabled gene delivery could treat DMD disease *in vivo*, we packaged our two therapeutic strategies into AAV vectors optimized for different delivery routes. For local intramuscular delivery to TA, REJ constructs were packaged with the AAV8 capsid. For systemic applications requiring broad biodistribution, we utilized the AAV-myo vector engineered for enhanced myotropic targeting across cardiac, skeletal, and diaphragm muscles^35^. We tested REJ-Abe8e gene editor in B6Ros.Cg-Dmd.mdx−4Cv/J (B6.mdx) mice and REJ-Dp253 gene replacement therapy in the more severe DBA/2-Dmd.mdx/J (D2.mdx) model using both local intramuscular and systemic delivery approaches (Figure 2A, E)^34,36–39^.

In treated TA muscle, REJ-Abe8e successfully restored full-length dystrophin levels to 19% of wildtype on average, with some samples reaching up to 66% (Figure 2C, D). REJ-Dp253 was robustly detected within muscle samples at an average of 72% of wildtype levels, with some samples reaching 195% (Figure 3A, B). Notably, the untreated TA muscles exhibited prominent plaques of damaged tissue, while the REJ-Dp253 treated muscles resembled wildtype tissue (Figure 3C).

**Figure 3.**
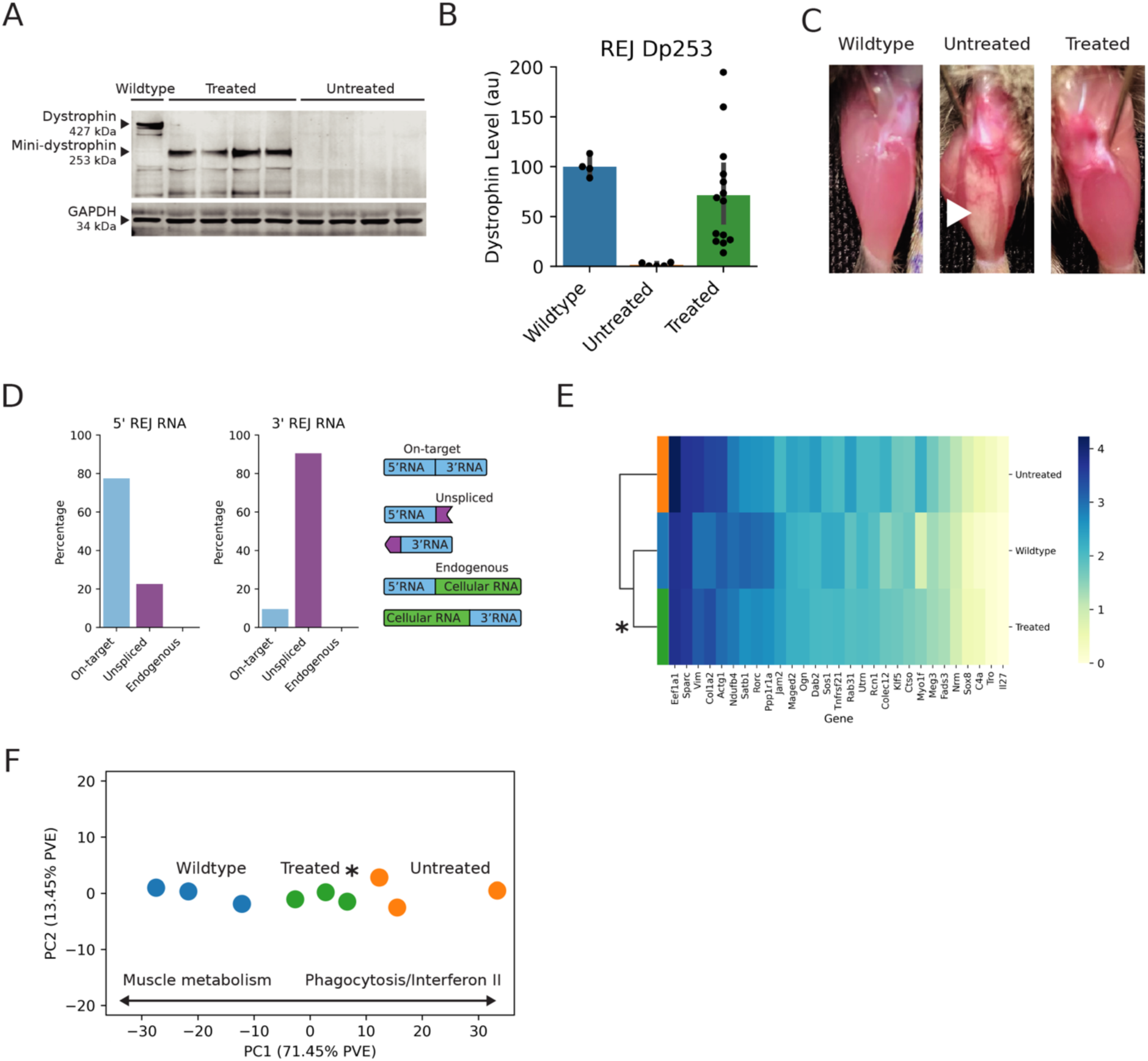
REJ-Dp253 restores dystrophin and improves the dystrophic transcriptome. A) DMD mice were treated with dual AAVs encoding Dp253, resulting in robust expression. B) Dual AAV Dp253 treatment resulted in 71.6% of wildtype dystrophin levels compared to 1.7% in untreated mice (p=0.0003). C) DMD mice develop white fibrotic patches (white arrowhead) visible on exposed muscle tissue. In contrast, Dp253 treated TA muscles remain healthy and resemble those of wildtype mice. D) RNA-Seq of transduced tibialis anterior muscle demonstrates the high precision of REJ trans-splicing. For each of the 5’ and 3’ RNAs, 78% and 9% of RNA were spliced on-target. Off-target splicing to endogenous transcripts was not detected (0.00%) with a statistical upper bound of <0.2% (95% CI 0.00-0.190%), confirming the high specificity of the REJ system. E) Expression of known DMD associated genes, spanning structural, transcriptional, metabolic, immune, and signaling proteins showed improvement in 24 of 29 genes (p<0.0001). F) A primary disease-associated axis was defined by performing principal component analysis (PCA) and gene ontology annotation with wildtype and untreated controls, revealing that muscle metabolism and phagocytosis/interferon II were major drivers of disease. Projecting the treated mice onto this axis revealed that their transcriptomic profile shifted significantly toward the wildtype cluster (p=0.0468)

REJ vectors expressing reporter proteins have an extremely low probability of off-target splicing to endogenous transcripts (Bachmann et al., companion manuscript, bioRxiv 2026). To investigate whether REJ-Dp253 and REJ-Abe8e were prone to off-target splicing with cellular transcripts we performed deep RNA sequencing. We found that 78% of the total 5’-REJ-Dp253 RNA and 9% of the total 3’-REJ-Dp253 RNA were spliced together to make full-length mRNA (Figure 3D). Although the dual AAV vectors encoding Dp253 were mixed at 1:1 titer ratios, the difference in splicing efficiency of the 5’-RNA segment relative to the 3’-RNA was likely due to higher expression or stability of the 3’ segment. Despite the relatively large amount of unspliced 3’-RNA in muscle we did not identify trans-splicing to endogenous RNA (Figure 3D), leading to an off-target splicing rate that is not statistically distinguishable from 0% (95% CI 0.000–0.190%). Comparable results were seen with the REJ-Abe8e treated mice (Figure S1C). These results further establish the high specificity of REJ.

We characterized the transcriptomic profile of muscle tissue from REJ-Abe8e and REJ-Dp253 treated mice to assess the overall physiology of cells. First, we evaluated expression of 29 diverse disease-associated genes, including the dystrophin homologue Utrn, structural cytoskeletal protein Vim, and the inflammatory cytokine Il-27^40–42^. In both treatment groups, the number of disease genes with improved expression was significantly shifted toward wildtype muscle levels (Figure 3E, Figure S1A). To extend this analysis and identify broader disease-specific gene expression patterns, we performed principal component analysis of all transcripts in muscle. This analysis clearly distinguished wildtype from untreated DMD samples (Figure 3F, Figure S1B). As expected, wildtype muscle was enriched for genes involved in muscle contraction and metabolism, while DMD muscle showed increased inflammation and immune response genes. Importantly, with both REJ-Abe8e or REJ-Dp253, the overall gene expression profile in treated muscle shifted significantly toward the wildtype pattern.

### Treatments restore physiologic muscle function and voluntary activity

To evaluate the functional efficacy of REJ-mediated dystrophin restoration, we performed comprehensive assessments spanning force generation and integrated whole-body activity^43^. In TA muscles, both REJ-Abe8e and REJ-Dp253 treatments significantly enhanced contractile function. REJ-Abe8e resulted in 46.6% specific force rescue compared to untreated controls (177.0 vs 128.2 mN/mm², p=0.0164), while REJ-Dp253 achieved a 62.9% rescue (198.3 vs 153.8 mN/mm², p=0.0013), with the latter approaching wildtype levels (224.5 mN/mm², p=0.0762) (Figure 4A, B, left panels).

**Figure 4.**
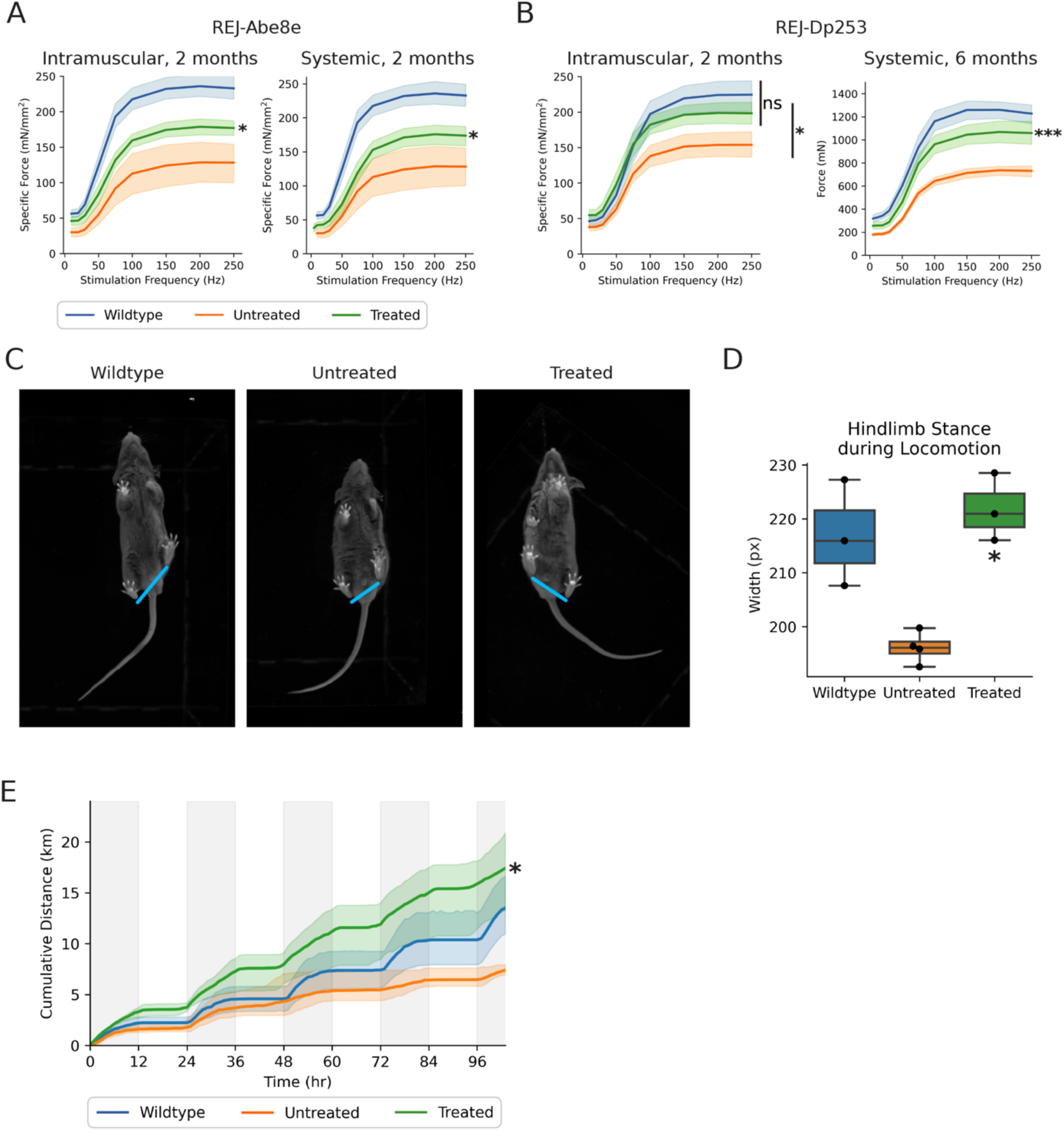
REJ treatment improves contractile force, gait, and voluntary activity. A) Mice treated with Abe8e intramuscularly (left) produced higher specific force (sPo) than untreated mice (177.0 vs 128.2 mN/mm^2^, p=0.0164) but not to the level of wildtype mice (232.9 mN/mm^2^) at 250 Hz. Mice treated systemically (right) with using a myotropic AAV vector exhibited higher specific force than untreated mice (173.8 vs 128.2 mN/mm^2^, p=0.024). Mice were evaluated at 2 months of age with the same wildtype and untreated controls. n=8-15 per group. B) Mice treated with Dp253 intramuscularly (left) produced higher specific force than untreated mice (198.3 vs 153.8 mN/mm^2^, p=0.0013) and were not significantly different than wildtype mice at maximum stimulation (224.5 mN/mm^2^, p=0.0762) at 2 months. In a systemically treated aged cohort (6 months, right), mice exhibited improved Po force generation compared to untreated mice (1058.7 mN vs 731.6 mN, p<0.0001), but not to wildtype levels (1227.5 mN, p<0.0001). n=9-15 per group. C) Hindlimb stance width was measured during free locomotion as an indicator of gait stability in mice treated systemically with REJ-Dp253. Representative images of paw placement, captured from below and tracked by deep-learning pose estimation, demonstrate the narrowed hindlimb stance and gait width of untreated mice, which is absent in wildtype and treated animals (blue). D) Untreated mice exhibited a significantly narrower stance width than wildtype controls (p<0.001). REJ-Dp253 treatment significantly rescued this phenotype (p=0.009), resulting in a stance width statistically indistinguishable from wildtype mice (p=0.51). p-values were calculated using Student’s t-test, n = 3-4 per group. E) On the voluntary running wheel assay, treated mice ran significantly further than untreated mice (17.4 km vs 7.4 km, p=0.0443), similar to wildtype mice (13.5 km, p=0.2305), demonstrating recovery of sustained and self-motivated exercise capacity. Night and day cycles are shaded gray and white respectively. n=3-5 per group.

Systemic delivery via myotropic AAV vectors maintained therapeutic efficacy, with REJ-Abe8e producing comparable force improvements (173.8 vs 128.2 mN/mm², p=0.024) (Figure 4A, right). Remarkably, in a cohort treated systemically from p14 and evaluated at 6 months, representing prolonged therapeutic exposure during critical developmental and degenerative phases, REJ-Dp253 treatment yielded a 66.0% absolute force rescue (1058.7 vs 731.6 mN, p<0.0001), demonstrating durable therapeutic benefit over extended treatment periods (Figure 4B, right).

Beyond force generation, this treatment prevented disease-associated wasting and restored treated mice body weight to wildtype levels (Figure S2B). To quantify locomotor mechanics, we employed deep-learning based pose estimation using Blackbox Bio and DeepLabCut to map limb positioning during ambulation. This analysis demonstrated that treatment corrected the pathological narrowing of hindlimb stance width, restoring a stable wildtype gait profile (Figure 4C, D).

Together, these improvements translated to profound restoration in voluntary physical activity. Mice systemically treated with REJ-Dp253 exhibited a 2.4-fold increase in cumulative running wheel distance compared to untreated controls (17.4 vs 7.4 km, p=0.0443), achieving performance not distinguishable from wildtype animals (Figure 4E). This restoration of voluntary exercise capacity is particularly striking given that sustained wheel running demands coordinated function across multiple organ systems affected in DMD, including cardiac muscle for circulation, diaphragm for respiration, and skeletal muscles for locomotion. The normalization of this complex, self-motivated behavior provides compelling evidence for the systemic therapeutic efficacy of REJ-mediated dystrophin expression.

### Dystrophin is restored across widespread muscle groups

Sections of the TA muscle of DMD mice treated intramuscularly with REJ-Abe8e or REJ-Dp253 were stained to label dystrophin, laminin-α2, and nuclei (Hoechst). Dystrophin expression was localized to the laminin-α2+ membrane of muscle fibers of both REJ-Abe8e and REJ-Dp253 treated mice, although more cells appeared to express dystrophin in animals treated with REJ-Dp253 (Figure 5A, B). Moreover, systemic delivery of AAV-myo REJ-Abe8e and REJ-Dp253 led to widespread dystrophin expression across muscle groups (Figure 5C, D; Figure S2A). Importantly, diaphragm and cardiac myofibers displayed robust dystrophin expression, which are two particularly sensitive muscle types in DMD patients.

**Figure 5.**
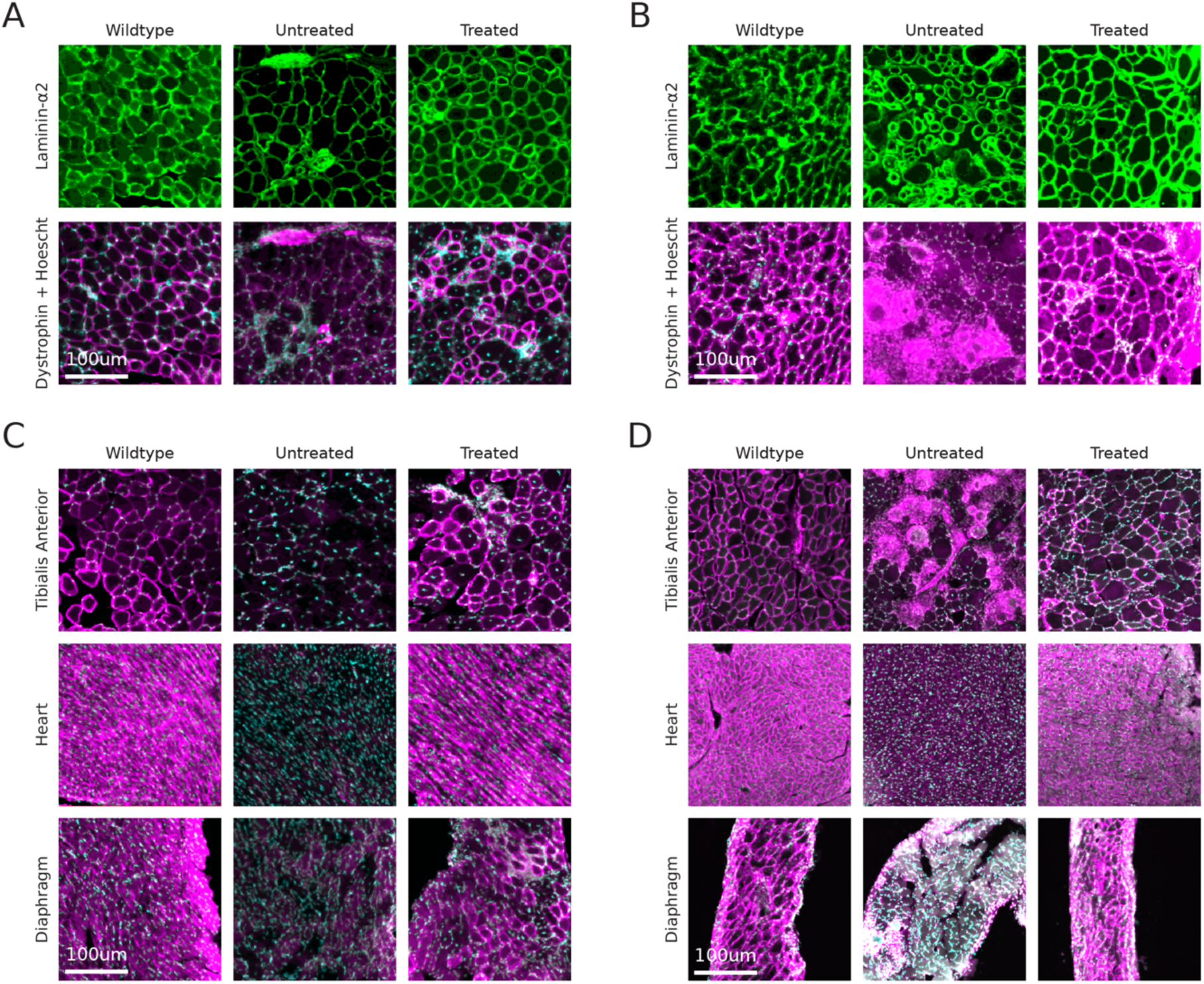
REJ-Abe8e and REJ-Dp253 AAV treatments enable widespread dystrophin expression. A, B) Abe8e (A) and Dp253 (B) delivered by direct intramuscular injection demonstrate proper localization of the restored dystrophin protein to the myofiber membrane (laminin-ɑ2 in green, Hoechst nuclear stain in blue, dystrophin in red). C, D) Abe8e (C) and Dp253 (D) delivered by systemic treatment with a myotropic AAV result in robust expression of dystrophin protein across all muscles analyzed, including skeletal, cardiac, and diaphragm.

### Whole-muscle single cell analysis using machine-learning and computer vision

To accurately quantify the effect of our treatments on the pathologic signatures of dystrophic muscle in DMD, we developed a high throughput image analysis pipeline to analyze tissue sections labeled with laminin-α2 (myofiber membrane), Hoechst (nuclei), and dystrophin (restored protein) (Figure 6A). The imaging pipeline we developed uses advanced computer vision and machine learning to rapidly quantify immunohistochemical features at the level of each individual cell, allowing for a high-throughput and statistically robust evaluation of muscle pathology (Figure S3)^44–46^. Using this workflow, all ∼2,500 myofibers within the cross-section of each TA muscle were computationally segmented (i.e. identified as individual cells) and phenotypically characterized using machine-learning and computer vision to define the subcellular location of nuclei, muscle fiber diameter, dystrophin-status, dystrophin expression level, and presence of infiltration from inflammatory cells. In total, over 120,000 myofibers were segmented and characterized, allowing for highly sensitive statistical analysis (Figure S3). Each mouse genetic background giving rise to DMD displayed unique pathological phenotypes. Consistent with previous reports, the automated high throughput computer vision characterization detected these cellular alterations within each DMD model with robust statistical differences compared to wildtype controls (Figure 6B-E).

**Figure 6.**
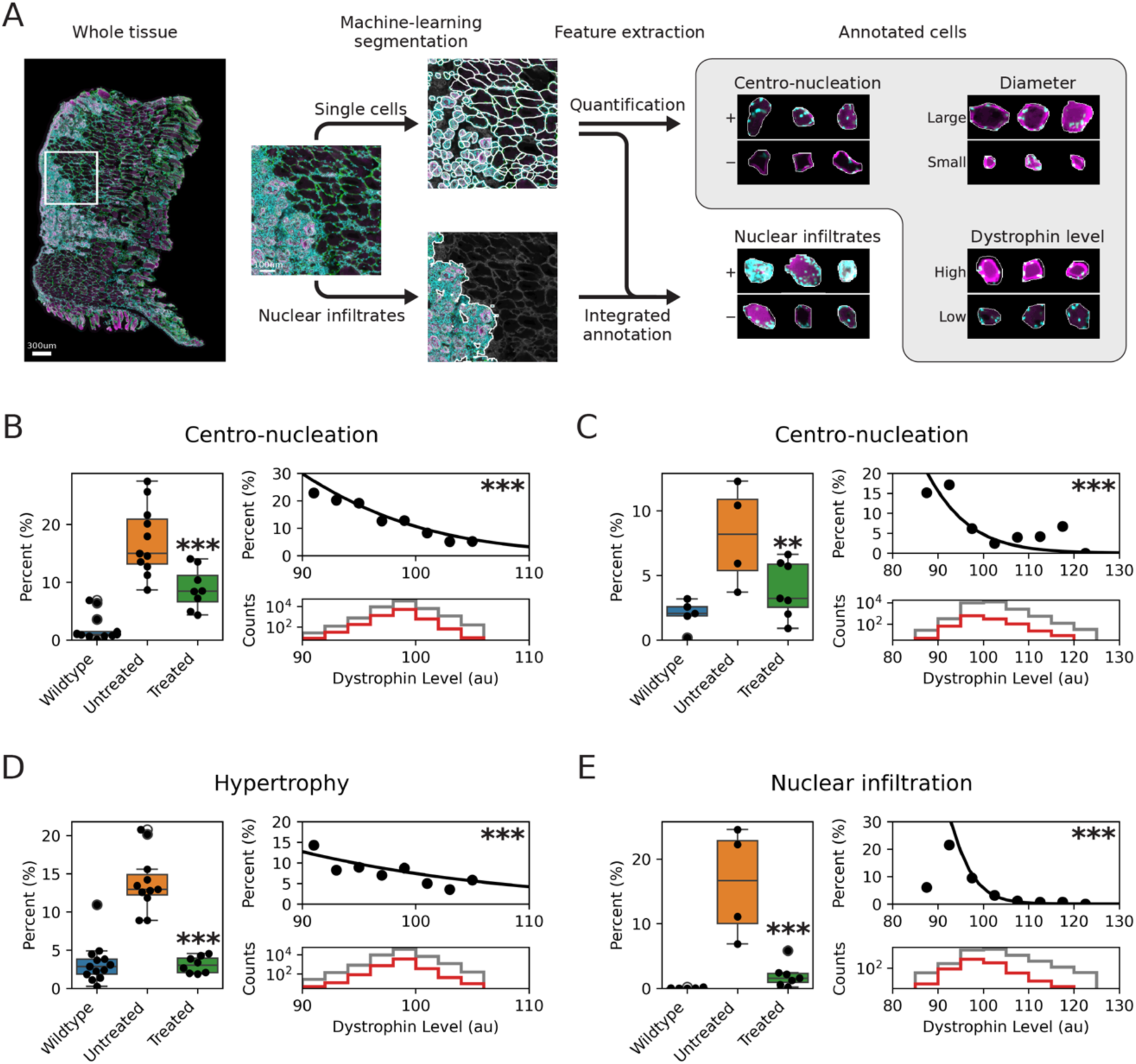
Advanced histopathological evaluation of muscle restoration using computer vision. A) Development of an advanced computer vision and machine learning histopathology pipeline enables robust statistical phenotyping across hundreds of thousands of individual myofibers. Using a deep-learning model, individual myofibers are segmented. Phenotypes are quantified by computer vision (gray) and machine learning implementations. B, D) Abe8e-treated mice exhibited improvements in centro-nucleation (wildtype 1.7%, untreated 15.7%, treated 8.8%, p<0.0001 treated vs. untreated). Hypertrophy was improved to wildtype levels (wildtype 2.9%, untreated 12.9%, treated 3.0%, p<0.0001 treated vs. untreated). Both phenotypes were negatively associated with dystrophin level (both p<0.0001, top right). Distribution of myofiber dystrophin signal intensity across treated (gray) and untreated (red) mice is shown. Data from n=8-11 mice per group with >30,000 myofibers per group. C, E) Dp253 treated mice showed improved centro-nucleation (wildtype 2.1%, untreated 7.2%, treated 3.25%, p=0.0125 treated vs. untreated) and nuclear infiltration (wildtype 0.06%, untreated 14.2%, treated 2.1%, p<0.0001 treated vs. untreated). Both phenotypes were negatively associated with dystrophin level (both p<0.0001, top right). Distribution of myofiber dystrophin signal intensity across treated (gray) and untreated (red) mice is shown. Data from n=4-7 mice per group with >7,500 myofibers per group.

In the gene-editing mouse model, the untreated mice exhibited an increased proportion of both centrally nucleated and hypertrophic myofibers. Both of these dystrophic features were significantly improved following REJ-Abe8e treatment (Figure 6B, D). In the gene-replacement mouse model, the untreated mice had an increased proportion of centrally nucleated myofibers and nuclear-infiltrates among the myofibers. RNA sequencing of the dystrophic tissue revealed a predominant immune signature of macrophages (∼43%) and dendritic cells (∼31%) consistent with these nuclear infiltrates representing active inflammatory infiltrates. Likewise, these pathological features were significantly reduced in the REJ-Dp253 treated mice (Figure 6C, E). The ability to examine the phenotypes of a large number of individual myofibers revealed a direct relationship between cellular dystrophin protein level and disease phenotype reduction (Figure 6B-E).

### Machine-learning reveals improved myofiber health in REJ-Abe8e

While we visually identified a significant proportion of myofibers expressing dystrophin in the gene-editing mouse model, amplicon sequencing indicated only 4% of the muscle cell genomes were edited by REJ-Abe8e. This was attributed to poly-nucleation of muscle fibers, where only a fraction of nuclei need to undergo *dmd* editing to restore dystrophin expression within a myofiber. To more accurately define the proportion of cells expressing dystrophin following gene therapy treatment, we trained a machine-learning classifier on 2,300 manually labeled myofibers images using an iterative active-learning approach validated by 20-fold cross-validation (Figure 7A, Figure S4)^47^. This classifier was then used to examine ∼30,000 muscle fibers following treatment with REJ-Abe8e with a comparable number of myofibers from wildtype and untreated controls. We found that 45.5% of myofibers in mice treated with REJ-Abe8e expressed dystrophin (Figure 7B). It has been shown previously that the *dmd* gene mutation reverts to wildtype in a low percentage of muscle fibers from B6.mdx^4cv^ mice^48^. Consistent with this reversion process, the classifier detected a small number of dystrophin-expressing cells in untreated animals (Figure 7B).

**Figure 7.**
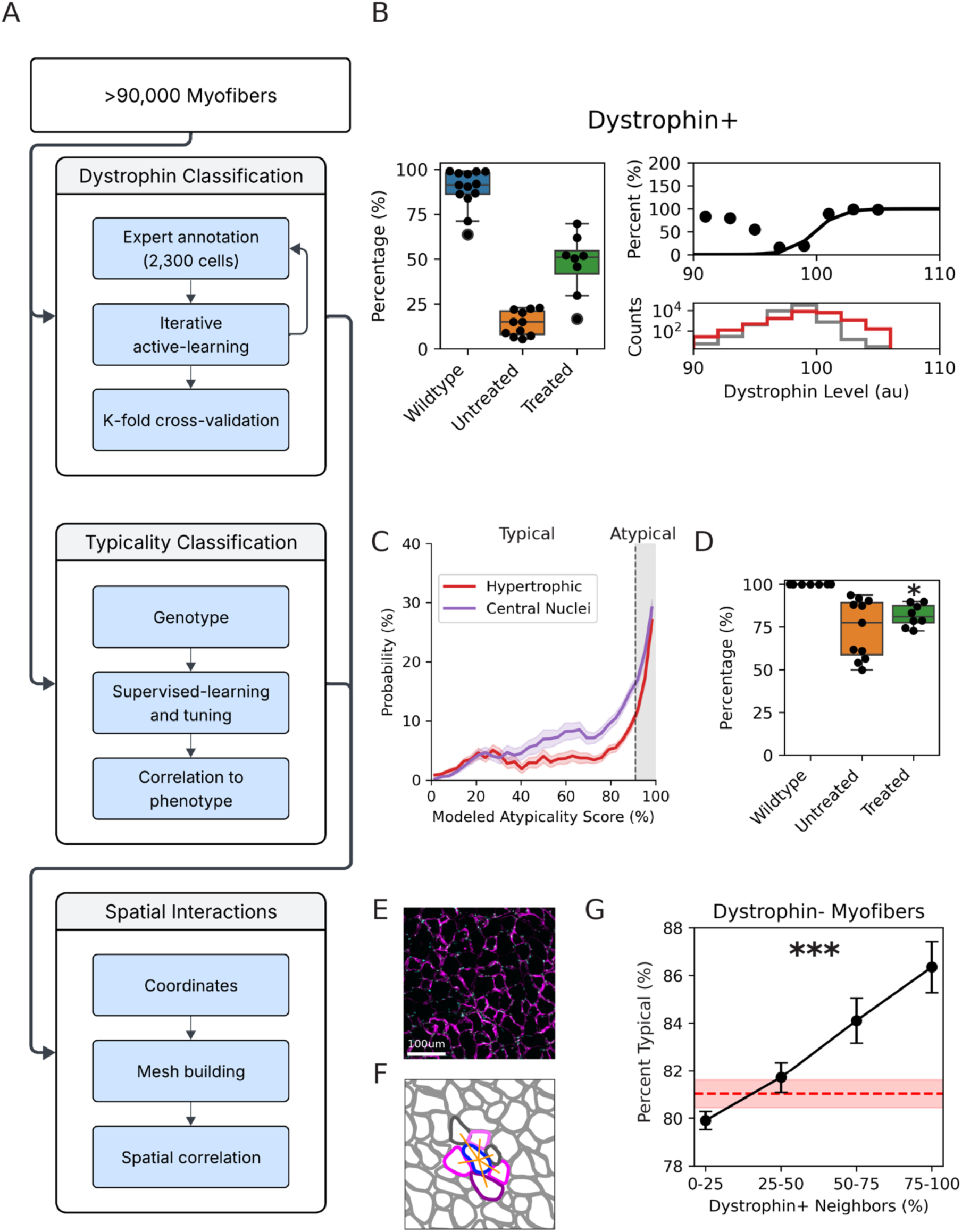
Machine-learning quantifies complex phenotypes and reveals spatial cooperativity. A) A suite of machine learning-based analysis frameworks was developed to assess complex attributes across >90,000 myofibers in Abe8e-treated mice. The quantification of dystrophin positivity, myofiber atypicality, and spatial interactions each required unique considerations of input data, model structure, and validation. B) A dystrophin+/– machine learning model was trained and validated on a 2,300 myofiber expert-annotated dataset (ROC AUC 0.87±0.03) and applied to the entire Abe8e dataset. This revealed that 45.5% of treated fibers expressed dystrophin compared to 13% revertant fibers in untreated and 89.2% in wildtype (p<0.0001, treated vs. untreated). C) A myofiber atypicality classifier was created using a genotype-supervised training approach. Myofiber atypicality was associated with centro-nucleation and hypertrophy across all myofibers and each treatment conditions. D) The formation of typical fibers was increased in treated myofibers compared to untreated fibers (81.9% vs 70.0%, p=0.0135). E) Female *Dmd* carriers do not exhibit disease phenotypes despite having X inactivation leading to mosaic expression of dystrophin (red). Laminin-α2 is shown in green. F) Spatial mesh construction using myofiber coordinates enabled precise neighborhood analysis for every myofiber within the muscle architecture. For each myofiber (blue), proximity (orange) to adjacent myofibers which have may have varying dystrophin expression statuses (purple). G) Spatial network analysis revealed a positive relationship between the percentage of dystrophin+ neighbors and typicality probability in dystrophin-fibers (rho=0.0405, p<0.0001). This correlation was not present in edited, dystrophin+ fibers (rho=0.01, p=0.2266). Null hypothesis and 95% CI in red.

To comprehensively assess therapeutic efficacy beyond conventional pathological markers, we developed a machine-learning approach to quantify the full spectrum of dystrophic alterations. We trained a classifier utilizing laminin-α2 membrane structure and nuclear organization patterns while excluding dystrophin signal (Figure 7C). This genotype-supervised approach generated a continuous atypicality score that captured the probabilistic health state of each myofiber, ranging from typical (healthy) to atypical (dystrophic) phenotypes (Figure 7C). Validation of this classifier against established disease markers confirmed its biological relevance, revealing strong correlations with both centro-nucleation (1.25 odds, p<0.0001) and fiber hypertrophy (1.35 odds, p<0.0001) across all experimental groups. Importantly, these correlations persisted within individual treatment arms, demonstrating that the classifier captured pathological signatures rather than batch differences. The typical/atypical score threshold was computed using the Youden index to maximize the sensitivity and specificity of detecting the previously characterized phenotypes.

Application of this model to the complete dataset revealed that 70.0% of untreated DMD myofibers exhibited typical morphology, whereas REJ-Abe8e treatment significantly improved fiber morphology, with 81.9% of myofibers classified as typical (p=0.0135, Figure 7D). This machine-learning approach thus provides a unified, sensitive, and unbiased single metric for therapeutic assessment that integrates both known and unknown visual pathological features beyond those detectable by conventional analyses.

### Dystrophin-positive fibers protect neighboring untreated cells

Muscle fibers are organized into fascicles surrounded by perimysium that represent functional units for muscle contraction^49^. Interestingly, female carriers of DMD mutations, which exhibit mosaic dystrophin expression due to X-chromosome inactivation, typically remain asymptomatic despite having a mixture of high- and low-dystrophin level fibers (Figure 7E)^1,2^. This raised the possibility that interactions between muscle fibers compensate for variable dystrophin expression. The chimerism we observed in REJ-Abe8e treated muscles (Figure 5A; 7B) and the single cell resolution in our computer vision framework allowed us to investigate the phenotype of dystrophin-negative fibers in the proximity of dystrophin-positive fibers. We implemented spatial network analysis to identify physical neighbors within the muscle architecture and integrated this information with the dystrophin-status and typicality of each cell (Figure 7F). This approach revealed that unedited myofibers had significantly improved myofiber typicality as the proportion of dystrophin+ neighbors increased (Figure 7G). This finding suggests that unedited myofibers in mice treated with REJ-Abe8e may be protected from damage if surrounded by dystrophin+ neighbors.

### Therapeutic full-length dystrophin delivery via triple REJ-AAV therapy

While using base-editors such as Abe8e to restore full-length dystrophin may be a viable treatment for DMD, clinical considerations remain, including off-target editing, immunogenicity of the effector, and gene mutation specificity^50,51^. Likewise, gene-replacement with REJ-Dp253 demonstrated robust therapeutic efficacy, but lacks several functional domains present in native dystrophin that may be critical for complete physiologic restoration. The full-length 427 kDa dystrophin protein contains all structural and regulatory elements, including domains absent in even the most promising mini-dystrophin variants^1^. However, the 11 kb coding sequence exceeds the capacity of dual-AAV systems, necessitating development of an even more sophisticated delivery strategy. To address this challenge, we engineered a triple-vector REJ-Dp427 system employing two orthogonal sets of REJ RNA dimerization domains to promote ordered assembly of three RNA fragments (Figure 8A, Bachmann et al., companion manuscript, bioRxiv 2026).

**Figure 8.**
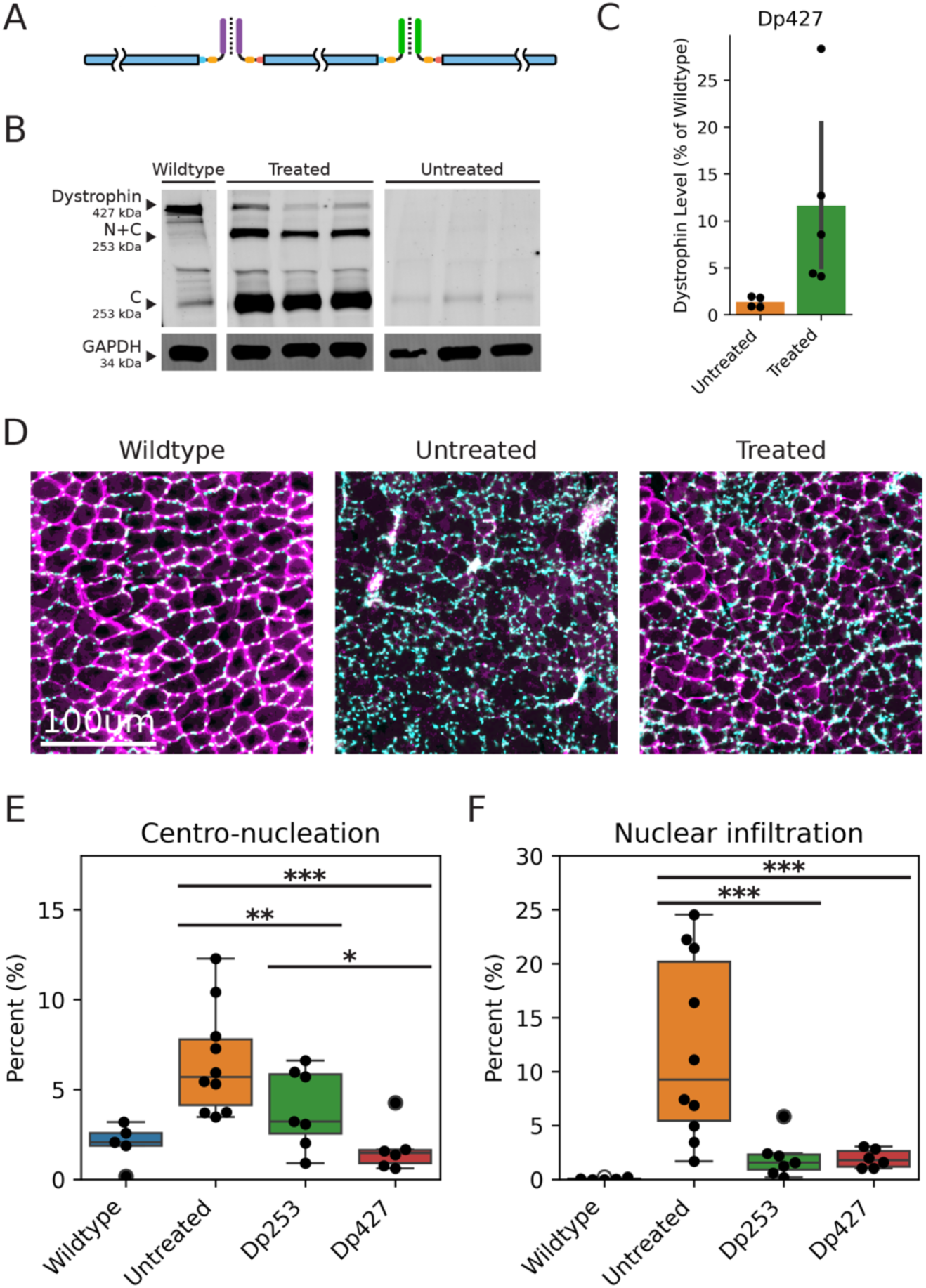
Triple-vector REJ system delivers therapeutic full-length native dystrophin. A) A triple AAV expression system carrying the native Dp427 was designed using two sets of linear dimerization domains. B, C) Mice treated intramuscularly expressed full-length dystrophin and achieved 11.6% of wildtype levels compared to 1.3% in untreated mice (p<0.05). Representative western blot shown. Lanes are not contiguous but derived from the same gel. D) Histology shows robust expression localized to the myofiber membrane and near saturation amongst all myofibers. E) In Dp427 treated mice, centro-nucleation is significantly improved compared to Dp253 and untreated mice (1.7% Dp427, 3.2% Dp253, p=0.0156). n=5-10 mice per group. F) In Dp427 treated mice, nuclear infiltration was reduced (p<0.01, 1.9% Dp427, 10.6% untreated) to comparable levels of Dp253 treated mice (2.1%, p=0.871).

Following intramuscular delivery of triple REJ-Dp427 vectors into D2.mdx mice, we achieved 11.6% of wildtype dystrophin levels (Figure 8B, C). Immunohistological analysis revealed near-complete myofiber saturation with proper sarcolemmal localization, confirming that REJ-mediated trans-splicing preserved the structural integrity of this massive protein and enabled correct integration into the dystrophin-sarcoglycan complex (Figure 8D). Remarkably, despite achieving lower overall expression than Dp253, full-length dystrophin provided significant therapeutic benefits in critical pathological metrics. Our sensitive computational histopathological framework revealed that centro-nucleation was significantly reduced to just 1.7% in Dp427-treated mice, representing a 47% relative improvement versus 3.2% centro-nucleation in Dp253-treated mice (p=0.0156) and approaching wildtype levels (Figure 8E). Nuclear infiltration decreased to 1.9% with Dp427, statistically matching the efficacy of Dp253 treatment (2.1%, p=0.871) and was dramatically improved compared to untreated controls (10.6%, p<0.01) (Figure 8F). Together, these results notably demonstrate the feasibility and potency of a triple-vector full-length dystrophin delivery system.

These results establish that REJ technology can scale beyond dual-vector limitations to deliver therapeutically relevant levels of full-length dystrophin. The superior histopathological outcomes achieved with native dystrophin, even at modest expression levels, convey the clinical importance of preserving functional domains and demonstrate that the REJ platform opens new therapeutic frontiers for genetic diseases caused by mutations in large, complex proteins previously considered inaccessible to AAV gene therapy.

## Discussion

This study demonstrates that RNA trans-splicing with REJ can overcome AAV cargo limits to express large therapeutic proteins in muscle. Dual-AAV systems produced functional levels of the 185 kDa base editor Abe8e and mini-dystrophin Dp253 at amounts comparable to wild-type dystrophin. Because both vectors must coinfect a cell, dual systems require 2.6-fold more virus than a single AAV to achieve similar coverage. Expression also depends on intracellular RNA::RNA interactions and efficient spliceosome reassembly, and studies with dual REJ vectors find it is ∼50% as efficient as a single-vector (Bachmann et al., companion manuscript, bioRxiv 2026). RNA sequencing suggests that balanced expression of both RNA segments is important because an underrepresented segment can become rate-limiting. Vector titration, together with optimization of capsid, promoter, dose, purification, and delivery route are important considerations for maximizing gene expression. Despite these variables, our findings using multiple dystrophin and base editor vectors establish that RNA trans-splicing is a reliable means of producing therapeutically relevant levels of proteins that cannot be encoded by a single AAV.

REJ-Dp253 treatment produced mini-dystrophin in nearly all myofibers, whereas REJ-Abe8e restored dystrophin in approximately half of muscle cells. Both approaches substantially improved muscle function, although the base-editor system may benefit from optimization. We did not measure Abe8e off-target DNA editing, an important issue under active investigation^11,33,52^. We did, however, assess unintended REJ-mediated splicing by RNA sequencing. Off-target joining to cellular transcripts was below detection at standard sequencing depth, indicating a very low error rate unlikely to meaningfully deplete endogenous RNAs.

Because full-length dystrophin Dp427 is more functional than truncated variants^5,12,53^, we also developed a triple-AAV REJ system. Two orthogonal RNA-dimerization pairs arranged three transcripts in the correct order and orientation, enabling two trans-splicing reactions and production of scarless, full-length dystrophin. This design is less efficient than the dual-vector platform and modeling predicts that 4.4-fold more virus than a single vector is required for triple coinfection. Nevertheless, REJ-Dp427 improved dystrophic muscle, consistent with clinical evidence that even low Dp427 levels can attenuate disease^54^. Thus, REJ can support three distinct therapeutic strategies, base editing, mini-dystrophin replacement, and full-length dystrophin replacement, although comparisons will be needed to determine their relative advantages.

Conventional histology, gene-expression, contractility, and behavioral assays all showed functional improvement after REJ-mediated dystrophin repair or replacement. To reduce bias and increase statistical power beyond that afforded by small sample sizes, we developed a high-throughput computational framework to analyze entire muscle cross-sections (>2,500 myofibers/sample). Deep-learning segmentation was used to analyze hypertrophy and central nucleation in over 120,000 myofibers, vastly exceeding manual methods. Random Forest models were trained, validated against expert annotations and biological ground truths, and classified nuclear infiltration, dystrophin positivity, and morphological “atypicality”. This platform provides a sensitive and unbiased method for evaluating therapeutic effects in complex disease models.

Remarkably, the depth and sensitivity of the computational platform developed here revealed non-cell-autonomous benefits of dystrophin expression in muscles. Dystrophin-negative fibers located near dystrophin-positive fibers were less likely to exhibit atypical morphology, suggesting that heterogenous expression of dystrophin among myofibers may still support muscle health. This observation could help explain why many female carriers of DMD mutations remain asymptomatic despite mosaic dystrophin expression caused by X-chromosome inactivation.

REJ compares favorably with other strategies for delivering large dystrophin constructs. Single-AAV micro-dystrophins such as Dp137 are clinically advanced^20,21,55^, but their truncation may limit function and efficacy^21^. Other multicomponent approaches include ribozyme-mediated RNA trans-ligation (StitchR)^13^, Cre-dependent DNA recombination (AAVLINK)^30^, and intein-mediated protein ligation^8,10,26^. Each has potential liabilities: StitchR has used viral doses near 1×10^15^ vg/kg, roughly tenfold higher than REJ and possibly above tolerated levels^56^; ribozyme cleavage products may affect cellular health; Cre can be mutagenic or toxic^57–61^; and inteins leave peptide scars, require abundant truncated intermediates, and may be immunogenic^8,10,26,27^. By contrast, REJ uses native splicing machinery, produces scarless proteins, functions efficiently, and showed no detectable off-target RNA interactions. Together with our computational assessment framework, these results support REJ as a versatile AAV platform with a clear translational path for DMD gene therapy.

## Methods

### Ethics statement

All animal procedures were approved by the Institutional Animal Care and Use Committee (IACUC) of the Salk Institute for Biological Studies (protocol 23-00020) and were carried out in accordance with the NIH Guide for the Care and Use of Laboratory Animals. Mice were group-housed (up to five per cage) in a specific-pathogen-free facility on a 12-h light/dark cycle with ad libitum access to standard chow and water, except during voluntary running-wheel experiments, for which animals were single-housed. Animals were randomly assigned to treatment and control groups. Male mice were used for all studies unless otherwise noted, because Duchenne muscular dystrophy is an X-linked disorder that predominantly affects males and the B6.mdx and D2.mdx models recapitulate disease in hemizygous males. At study endpoints, animals were euthanized by CO₂ asphyxiation followed by cervical dislocation.

### Animal models

Two dystrophin-deficient mouse lines were used. B6Ros.Cg-Dmd^mdx-4Cv^/J (B6.mdx; The Jackson Laboratory, stock no. 002378, RRID:IMSR_JAX:002378) mice were used for REJ-Abe8e gene-editing studies, because the mdx4cv allele carries the nonsense mutation targeted by the sgRNAs used here. D2.B10-Dmd^mdx^/J (D2.mdx; The Jackson Laboratory, stock no. 013141, RRID:IMSR_JAX:013141) mice, which develop a more severe dystrophic phenotype on the DBA/2 background, were used for REJ-Dp253 and REJ-Dp427 gene-replacement studies. Colonies of both lines were maintained in house from founders obtained from The Jackson Laboratory. Background-matched wild-type C57BL/6J (stock no. 000664, RRID:IMSR_JAX:000664) and DBA/2J (stock no. 000671, RRID:IMSR_JAX:000671) mice served as non-dystrophic controls for B6.mdx and D2.mdx experiments, respectively.

Intramuscular treatments were delivered to the tibialis anterior at postnatal day 28, and treated muscles were analysed at 2 months of age. Systemic treatments were delivered by retro-orbital injection at postnatal day 14; these cohorts were analysed at 2 months of age (REJ-Abe8e) or at 6 months of age (REJ-Dp253), with the 6-month cohort additionally undergoing voluntary running-wheel and body-weight monitoring. Group sizes for each experiment are reported in the corresponding figure legends.

### Plasmid design and construction

Plasmids were designed using SnapGene software and assembled using In-Fusion HD (Takara) or NEBuilder HiFi (NEB) kits. Constructs encoding split Abe8e, Dp253, or Dp427 were transformed into Stellar Competent Cells (Takara) and produced under antibiotic selection. Final plasmids were purified for transfection or viral packaging using QIAprep Mini (Qiagen), ZymoPURE Midi (Zymo), or PureLink Maxi (Invitrogen) kits. For structural visualization (Figure 1B), the micro-dystrophin protein structure was predicted using AlphaFold 2.

### Cell culture and in vitro transfection assays

HEK293T cells were maintained in DMEM (Gibco) supplemented to 10% FBS. Cells were seeded in 48-well plates and transfected at 80% confluency with 600 ng of each plasmid indicated using Lipofectamine 3000 according to manufacturer protocol. For REJ-Dp253 fragment suppression optimization, 5’ and 3’ REJ vectors were co-transfected at stoichiometric ratios as indicated. Cells were harvested 48 hours post-transfection for protein extraction. For base-editing validation, cells were co-transfected with a YFP reporter interrupted by the DMD^mdx.4cv^ nonsense mutation and imaged using an Olympus FV3000-RS microscope.

### AAV vector production and purification

Recombinant AAV was produced in HEK293T cells via triple transfection with Rep/Cap (AAV8 or AAV-myo), Ad5 helper, and transfer plasmids by the Salk Viral Vector core facility. Viral particles were harvested 72 hours post-transfection by three freeze-thaw cycles and Benzonase treatment, followed by precipitation with PEG 8000. Vectors were purified using either iodixanol gradients (15-60% step) or sequential CsCl density gradients (1.3-1.5 g/mL). Purified fractions were buffer-exchanged into PBS with 5% sorbitol, sterile-filtered, and titered by qPCR in triplicate.

### Stochastic viral infection modeling

Cellular transduction was modeled as a Poisson process where dystrophin-negative myofibers correspond to cells containing zero functional viral genomes (*k* = 0). The mean effective multiplicity of infection (*λ*) was calculated from the observed fraction of negative cells as *P*(*X* = 0) = *e*^-*λ*^. To relate physical viral titer to biological activity, an “expressivity” constant *ε* was defined such that *λ* = *ε* · *D*, where *D* is the total viral dose in vg. The value of *ε* was determined experimentally via linear regression calculated from single-vector AAV-Dp153 dose-titration studies. For multi-vector systems requiring n components, transduction efficiency was explicitly modeled as the joint probability of independent co-infection with at least one functional copy of each component as 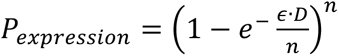.

### Intramuscular and systemic AAV administration

Mice were anesthetized using 2-3% isoflurane and assessed for depth of anesthesia via toe pinch. For intramuscular delivery, 10 µL of viral solution in AAV carrier buffer was injected into the proximal and distal segments of the TA muscle using pulled glass capillaries. For the single-vector titration analysis used to calibrate the stochastic infection model, AAV8-Dp153 vectors were administered at the specified doses diluted in 10 µL of AAV carrier buffer. For the therapeutic efficacy studies, total viral doses used were 1x10^11^ vg for REJ-Abe8e and 5x10^11^ vg for REJ-Dp253 and REJ-Dp427. Systemic delivery was performed in p14 mice via retro-orbital injection at a total dose of 1x10^14^ vg/kg and injection volume of 10 µL/g of body weight. Following injection, mice were monitored for the return of righting reflexes before being returned to their home cage.

### In situ contractile force measurement

*In situ* force generation of TA muscles was measured using an Aurora Scientific 1300A Whole Animal System by previously established protocol^43^. In brief, mice were anesthetized using 2-3% isoflurane, then tested for proper depth of anesthesia via a toe pinch. The distal TA tendon was detached and tied to the lever of a force transducer using silk suture. The optimal muscle fiber length (LO) was determined as length of the TA while held at 30 mN of tension. While held at LO, the muscle was stimulated with a range of frequencies and force generated was recorded. Specific muscle force values were obtained by normalizing the isometric tetanic force with cross sectional area. Cross sectional area was calculated using the following equation: CSA = (muscle mass, in gram)/[1.06g/cm3(0.6LO, in cm)].

### Voluntary running wheel, behavior, and weight monitoring

Systemically treated mice and controls were evaluated at 6 months of age. For voluntary exercise assessment, mice were single-housed in cages Colbourn running wheel cages (ACT-551SS) containing an 11 cm diameter running wheel equipped with mechanical switches (ActiMetrics) to record wheel turns. Mice were housed under a 12-hour dark-light cycle. Activity was recorded for 108 hours following a 96-hour acclimatization period.

For whole-body pose analysis, mice were recorded for 5 minutes during free locomotion in a Blackbox Bio apparatus. Pose estimation was performed using a pretrained DeepLabCut model and quantified in Python. Body weights were monitored at 4-week intervals starting from postnatal day 28.

### Protein extraction and western blot analysis

Total proteins were extracted from HEK293T cells by washing with PBS and resuspending in RIPA lysis buffer containing 4% protease inhibitor (Sigma) and 0.11% Benzonase. For muscle tissue, approximately 25 mg of sample was flash frozen in liquid nitrogen, pulverized for 3 cycles at 2000 rpm for 30s, treated with 200 µL of extraction buffer (RIPA with 10% SDS), and agitated for 10 minutes. Cell or muscle samples were then passed through an insulin syringe three times before being centrifuged for 10 minutes at 21,300 rcf. Supernatant was aliquoted and stored at - 20C.

### Genomic DNA extraction and sequencing

Genomic DNA was isolated from flash-frozen TA muscles (25 mg segments) using the DNeasy Blood and Tissue Kit (Qiagen) according to the manufacturer’s protocol. DNA amplification was performed using primers flanking the mdx.4cv mutation and purified by gel electrophoresis. Amplicons were sequenced on the Illumina platform (AmpliconEZ, Azenta) and analyzed using the CRISPResso2 package with default parameters.

### RNA extraction and sequencing

Total RNA was isolated from TA muscle tissue using the RNeasy Plus Universal Mini Kit (Qiagen). RNA integrity was assessed using an Agilent Bioanalyzer 4200. Library preparation and sequencing was performed by the Salk Next-Generation Sequencing Core. Libraries were prepared using the Illumina TruSeq Stranded mRNA kit with polyA enrichment. Sequencing was performed on an Illumina NovaSeq 6000SP with 150 bp paired-end reads at a target depth of 25 million reads per sample. Raw reads were quality controlled and trimmed using USEARCH12 and transcript abundance was quantified by mapping to the GRCm39 (mm39) reference using Salmon. Immune cell type fractions were estimated from Salmon-derived TPM counts using CIBERSORTx with the ImmuCC mouse muscle signature profile in relative mode. Differential gene expression analysis was performed using PyDESeq2. Principal component analysis (PCA) was performed using scikit-learn and gene ontology enrichment analysis using Enrichr. To quantify REJ splicing fidelity, custom Python scripts filtered for reads containing 25bp “anchor” sequences immediately upstream or downstream of the REJ splice junctions. The adjacent 15 bp sequences were extracted and mapped as on-target, unspliced, or endogenously spliced.

### Histology and immunofluorescence preparation

Mice were euthanized at the timepoints indicated and TA, triceps brachii, inferior cardiac ventricles, or diaphragm muscles were dissected, embedded in Tissue-Tek OCT compound, and frozen on dry ice. Sections were cut to 25 µm and captured on Superfrost Plus Microscope slides. For immunofluorescence, sections were fixed with ice-cold acetone for 10 minutes at -20C followed by blocking (10% NGS, 3% BSA in 1x PBS) for 30 minutes at room temperature. Primary antibodies targeting dystrophin, HA, and laminin-*α*2 were applied overnight at 4C. Slides were washed three times with PBS and incubated with secondary antibodies and Hoechst for 30 minutes. Slides were mounted using Mowiol and imaged using an Olympus FV3000-RS or Olympus VS-120 virtual slide scanning microscope at 4X.

### Automated myofiber segmentation and phenotyping

Whole-slide immunofluorescence images were processed to identify and characterize individual myofibers. Myofiber boundaries were segmented using the Cellpose 3 cyto3 deep-learning model on the laminin-*α*2 channel. For each segmented cell, morphological and intensity features were extracted using custom Python scripts utilizing the OpenCV library. The Feret diameter was calculated from binary cell masks to assess hypertrophy; fibers exceeding the mean diameter of wildtype controls by two standard deviations were classified as hypertrophic. Centro-nucleation was quantified by measuring Hoechst intensity within the cytoplasmic region, defined by eroding the cell mask using OpenCV morphological operations, and applying a quantile based intensity threshold.

### Machine learning classification frameworks

To quantify therapeutic efficacy beyond standard metrics, we developed Random Forest classifiers using the scikit-learn library to assess nuclear infiltration, dystrophin status, and myofiber atypicality. Nuclear infiltration was identified using a pixel-level Random Forest classifier trained on texture and intensity features extracted via scikit-image multiscale_basic_features. A Dystrophin positivity classifier was trained to identify dystrophin-positive fibers using dystrophin-channel intensity features. This model was optimized using an iterative active learning workflow starting with a seed set of 500 manually annotated myofibers. In subsequent rounds, the classifier identified 200 myofibers with low confident predictions for manual annotation and retraining with a total of 10 training cycles. To ensure dataset balance, these candidates were selected equally from the Wildtype, Untreated, and Treated groups. Finally, to generate an Atypicality score, we trained a separate Random Forest classifier to distinguish between myofibers from healthy and diseased mice based solely on structural markers, specifically Laminin-*α*2 and Hoechst, excluding the dystrophin channel to prevent data leakage. To mitigate technical variability between samples, input features were batch-corrected using Z-score normalization (scikit-learn StandardScaler) prior to classification. The model output provided a continuous probability score, which was binned into binary typical versus atypical categories by calculating an optimal probability threshold using the Youden Index to maximize the classifier’s agreement with the previously established pathological markers of hypertrophy and centro-nucleation.

### Spatial network analysis

To investigate non-cell-autonomous effects, we modeled the muscle tissue as a spatial graph. The geometric center of all segmented myofibers were used to construct a mesh network via Delaunay triangulation. Edges exceeding the 90th percentile of length distribution were pruned to remove connections across perimysial spaces or artifacts. This network allowed us to quantify the “neighborhood environment” for every cell. For each myofiber, we calculated the proportion of direct (1st-hop) neighbors that were dystrophin-positive. We then correlated this neighborhood fraction with the “Atypicality” score of dystrophin-negative cells to assess whether proximity to edited fibers conferred a protective morphological benefit.

### Statistics

Comparisons of means were performed as two-tailed Welch’s t-tests (unequal variances t-tests). Histological disease phenotypes were quantified and binned into binary classifications (see above). Comparisons were performed using Generalized Estimating Equations (GEE) with a Binomial family distribution to account for the hierarchical nesting of myofibers within individual muscles from three treatment groups, using the Python statsmodel package. Models defined the treatment group as the fixed effect and muscle identity as the grouping variable. The association between dystrophin level and each histological phenotype was modeled as a logistic regression across untreated and treated muscle fibers. Error bars and shaded area around means represent standard error unless otherwise noted. Confidence interval for endogenous off-target splicing was calculated as an exact binomial proportion. Relationship between dystrophin+ neighbors and dystrophin-myofiber health was calculated as a spearman correlation coefficient. Confidence intervals for proportions of endogenous trans-splicing were determined using the Clopper-Pearson exact method to establish a 95% confidence interval.

## Data availability

High-throughput sequencing data generated in this study have been deposited at the NCBI Sequence Read Archive (SRA) under NCBI BioProject accession PRJNA1480055 and are publicly available as of the date of publication. Plasmids generated in this study, including the split REJ-Abe8e, REJ-Dp253, and REJ-Dp427 constructs and associated control vectors, will be deposited to Addgene and made available as of the date of publication. Plasmids and other unique reagents that are not available through Addgene are available from the lead contact upon reasonable request. The B6.mdx (B6Ros.Cg-Dmd^mdx-4Cv^/J) and D2.mdx (D2.B10-Dmd^mdx^/J) strains are available from The Jackson Laboratory.

## Code availability

All original code, including the stochastic viral-infection model, the myofiber segmentation and feature-extraction pipeline, the machine-learning classification frameworks (dystrophin status, nuclear infiltration, and atypicality), the spatial network analysis, and the REJ splicing-fidelity quantification scripts, has been deposited on GitHub (https://github.com/ryanusahk/rej-dmd-histopathology-analysis) and is publicly available. Any additional information required to reanalyze the data reported in this paper is available from the lead contact upon request.

Use of generative AI and AI-assisted technologies in the writing process

During the preparation of this work, the authors used Claude (Anthropic) and ChatGPT (OpenAI) in order to proofread text for clarity. After using these tools, the authors reviewed and edited the content as needed and take full responsibility for the content of the published article.

## Acknowledgments

We thank the Salk Institute Next-Generation Sequencing Core, the Biophotonics Core, and the Gene Transfer, Targeting and Therapeutics (GT3) Core for their technical assistance and services. This work was supported by the Department of Defense (DOD), the Marshall Heritage Foundation, the G. Harold and Leila Y. Mathers Foundation, the Keck Foundation, and the Wu Tsai Human Performance Alliance. R.H.H. was supported by an F30 fellowship training grant from the National Institute of Child Health and Human Development (NICHD).

## Author Contributions

Conceptualization, R.H.H., C.E.W., G.M., S.K., L.C.B., and S.L.P.; Methodology, R.H.H., C.E.W., L.C.B., and S.L.P.; Software, R.H.H. and S.K.; Validation, R.H.H., C.E.W., and S.L.P.; Formal Analysis, R.H.H. and C.E.W.; Investigation, R.H.H., C.E.W., G.M., M.G., K.L., C.J.A., K.J.H., S.K., and L.C.B.; Resources, R.H.H., G.M., S.P., L.C.B., and S.L.P.; Data Curation, R.H.H. and C.E.W.; Writing – Original Draft, R.H.H., C.E.W., L.C.B., and S.L.P.; Writing – Review & Editing, R.H.H., C.E.W., G.M., L.C.B., and S.L.P.; Visualization, R.H.H.; Supervision, R.H.H., L.C.B., and S.L.P.; Project Administration, L.C.B. and S.L.P.; Funding Acquisition, R.H.H., S.K., L.C.B., and S.L.P.

## Supplemental Information

**Figure S1.**
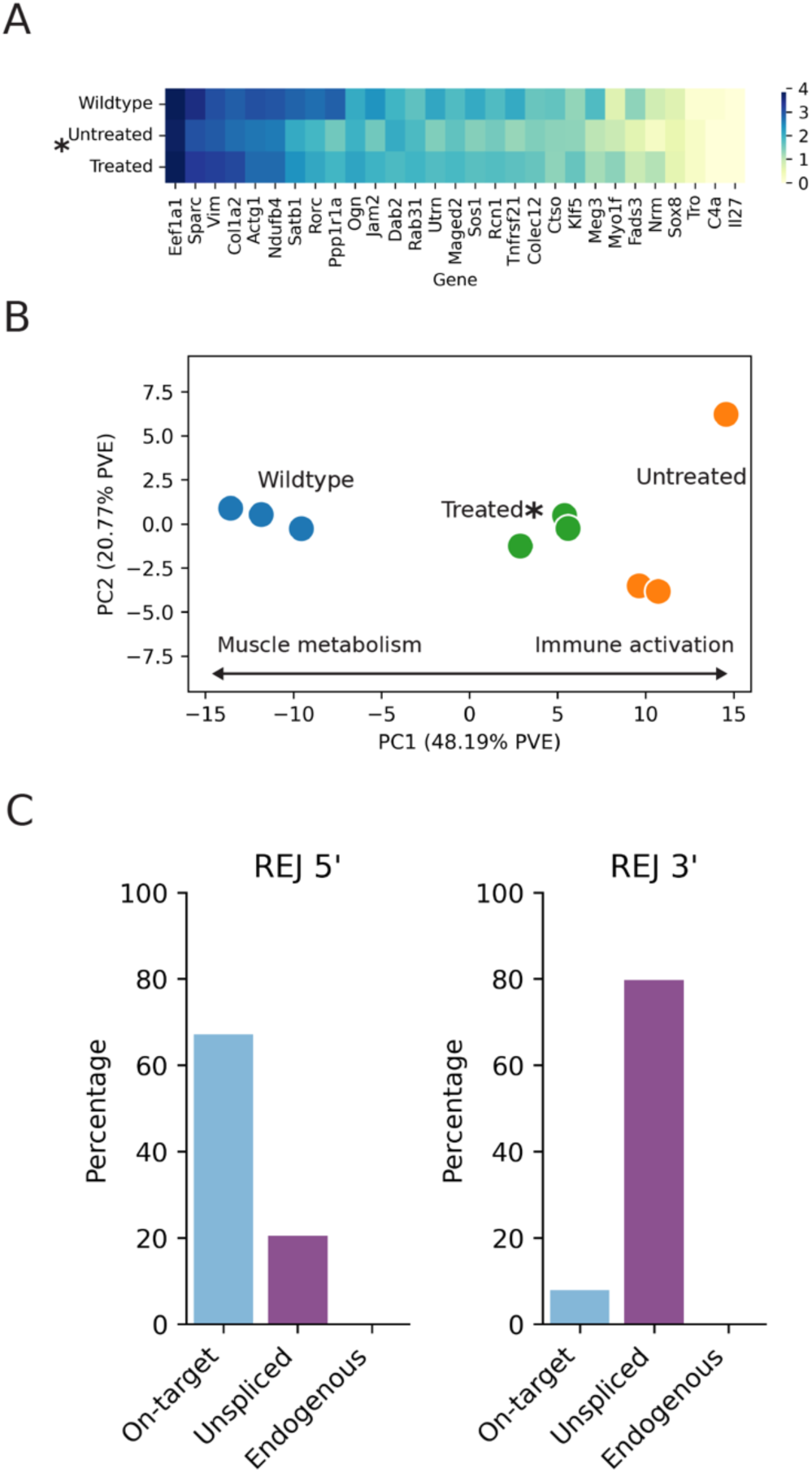
RNA analysis of Abe8e-treated mice. A) Expression of 29 diverse disease-associated genes showed significant restoration toward wildtype levels in REJ-Abe8e treated mice, with improvement observed in 21 of 29 genes (p=0.0242, n=3 per group). B) Principal component analysis (PCA) defined a primary disease axis between wildtype and untreated controls. The transcriptomic profile of REJ-Abe8e shifted significantly toward the wildtype cluster (p=0.0119). C) RNA-Seq quantification of splicing fidelity for the 5’ and 3’ REJ-Abe8e RNA. Off-target splicing to endogenous transcripts was not detected, with a statistical upper bound of <0.190% (95% CI 0.00-0.190%).

**Figure S2.**
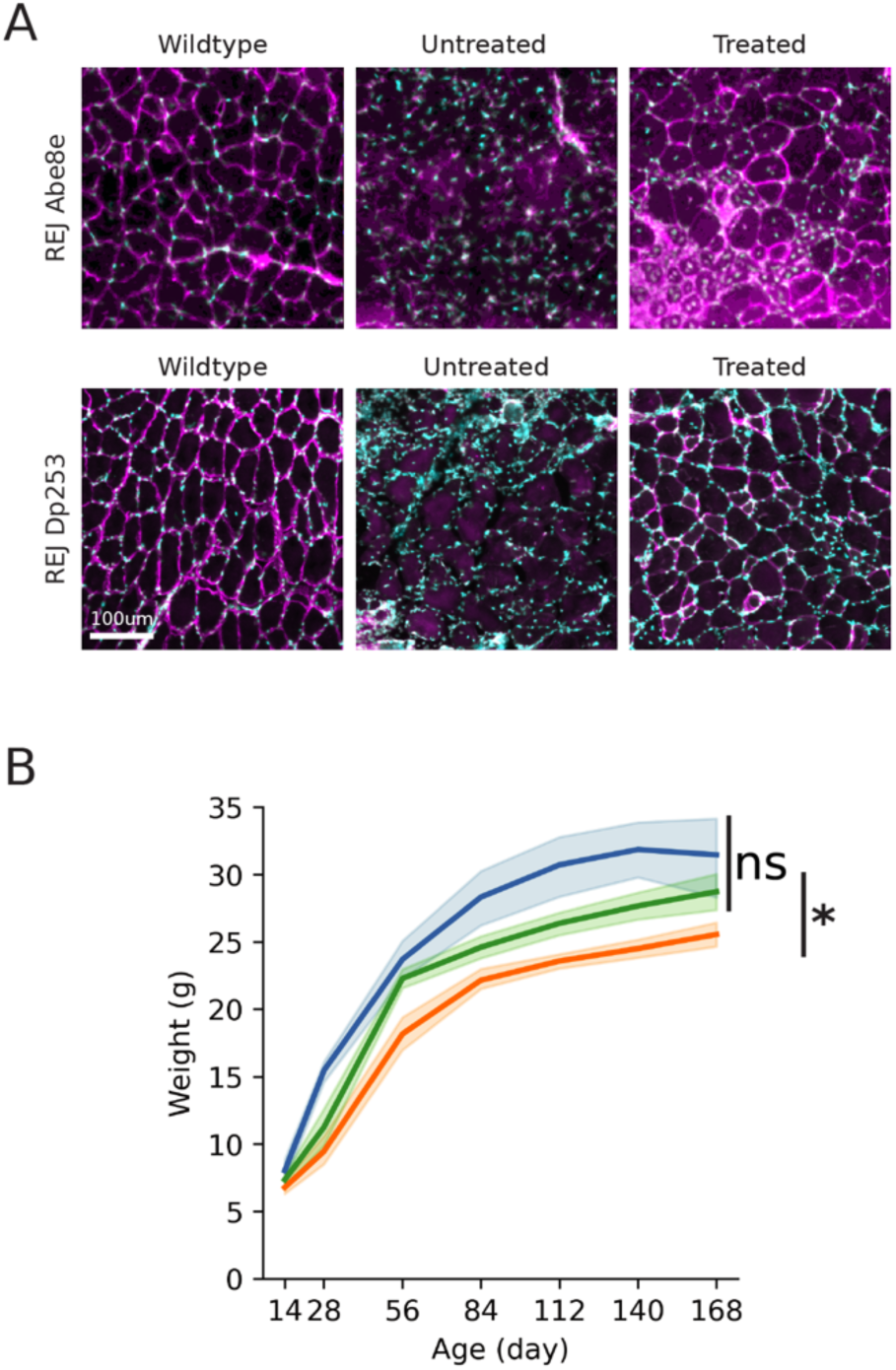
Systemic REJ treatment restores dystrophin to the triceps and prevents disease-associated wasting. A) Representative immunofluorescence images of triceps muscle sections following systemic treatment. The top row displays muscles treated with REJ-Abe8e, and the bottom row displays muscles treated with REJ-Dp253. Dystrophin (red) is restored in treated animals compared to untreated controls. Nuclei are stained with Hoechst (blue). B) Body weight monitoring demonstrates that Dp253-treated mice are protected from disease-associated wasting. Mice were treated at p14 and weighed from p28 at 4-week intervals. Treated mice were significantly heavier than untreated controls (p=0.0027) and were not statistically distinguishable from wildtype animals (p=0.1366). n=10-11 mice per group.

**Figure S3.**
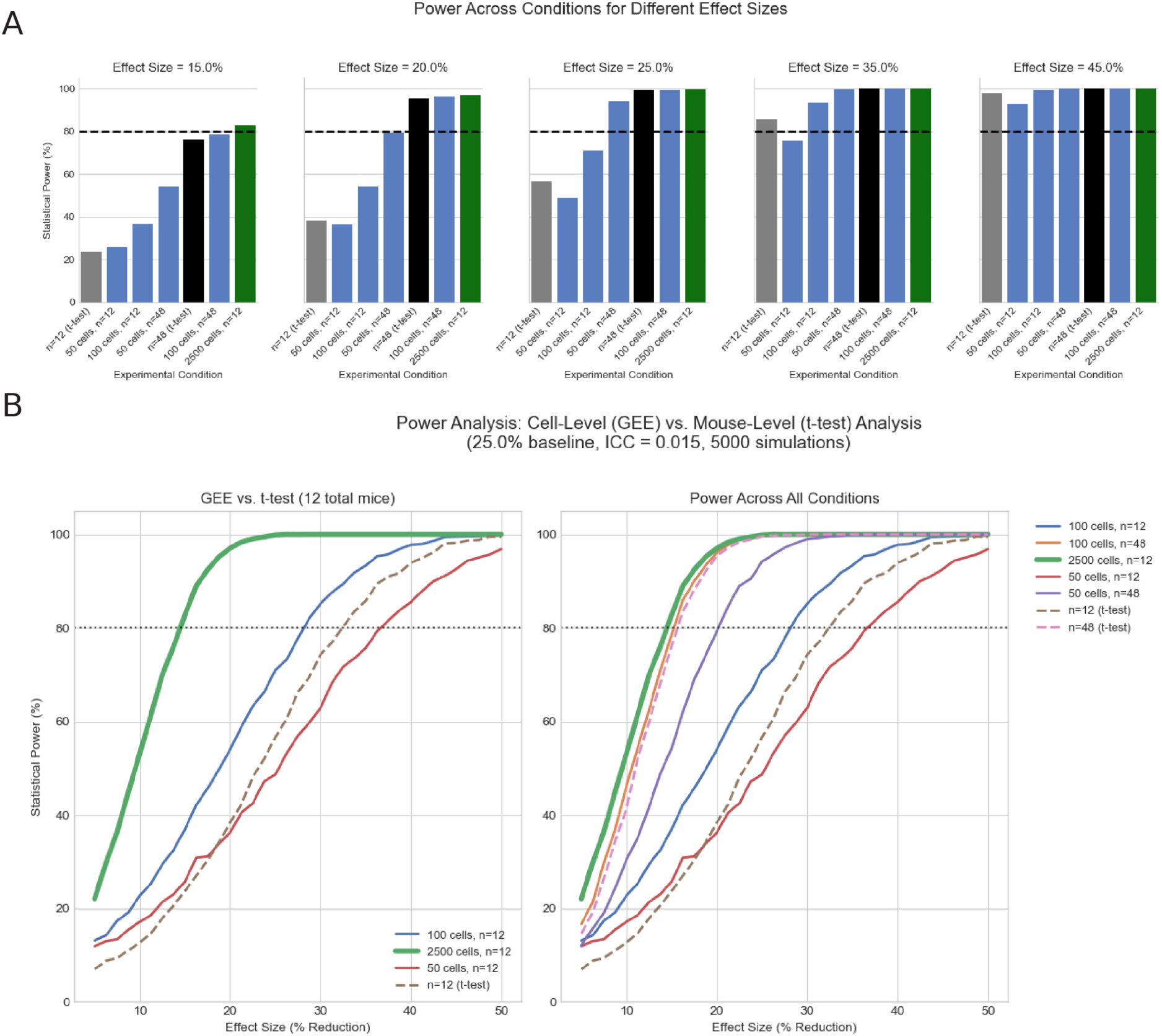
Statistical power analysis. A) Statistical power analysis comparing a standard mouse-level t-test (n=12, gray) against the cell-level Generalized Estimating Equations (GEE) model with increasing numbers of analyzed myofibers. The dashed line represents the standard 80% power threshold. For a subtle 15% therapeutic effect, the automated analysis of 2,500 cells yields 82.8% power, whereas the standard approach yields only 23.7%. B) Power curves plotting statistical power against effect size (% reduction in pathology). The green line represents the power achieved by the regime of the current study design (n=12 mice, ∼2,500 cells/mouse), which outperforms even a 48 mouse study with traditional averaging and t-tests. Power was estimated using 5,000 Monte Carlo simulations. The intracluster correlation (ICC) was set to 0.015, derived empirically from the variance observed in the experimental mouse samples.

**Figure S4.**
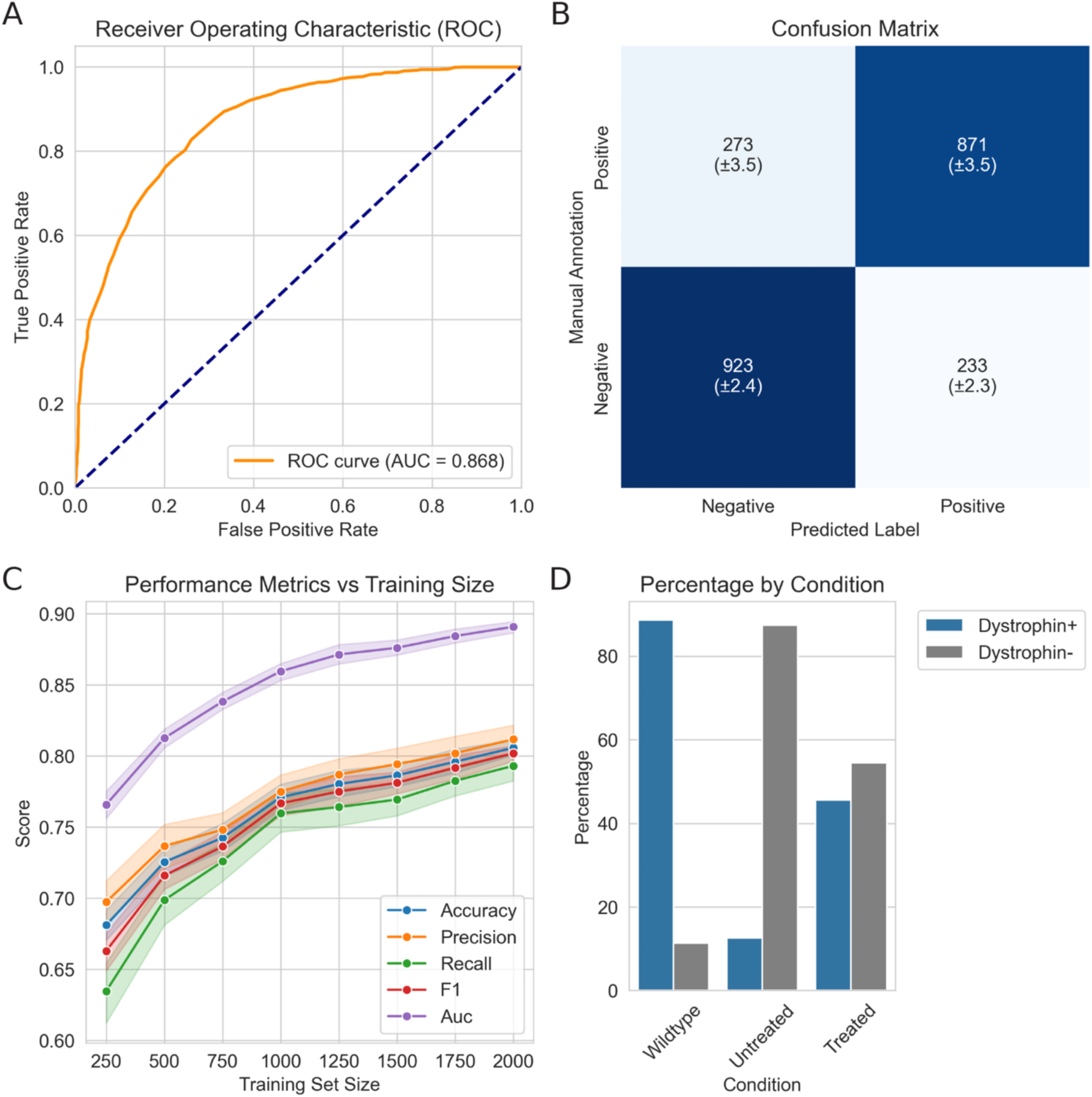
Dystrophin Positivity Machine Learning Model Validation. A) Technical validation of machine learning classifiers. Validation metrics for the dystrophin status classifier were calculated using 20-fold cross-validation on 2,300 expert-annotated myofibers (1,104 positive, 1,196 negative). The model demonstrated robust performance with a mean ROC AUC of 0.868 ± 0.03 and accuracy of 0.78 ± 0.04. B) Confusion matrix of pooled cross-validation predictions showing the distribution of classification results. C) Learning curves comparing model performance with training set size (250 to 2000 samples). Performance improves and stabilizes near the final training set size (2,300 samples). D) The percentage of positive and negative classified fibers across experimental conditions.

